# Betrayal is worse than loss during cooperation

**DOI:** 10.64898/2026.03.04.709582

**Authors:** Rumeng Tang, Jingbin Tan, Yi Gao, Chen Lin, Jing Gan, Xiaowei Ding, Dingguo Gao

## Abstract

Cooperative behavior refers to activities that benefit another individual or benefit both the person performing the behavior and the recipient. Although both “betrayal aversion” and “loss aversion” (the financial detriment incurred from betrayal) are established determinants of cooperative behavior, their relative potency remains undetermined. Here, we investigated these effects by integrating computational modeling and event-related potential (ERP) techniques. In two tasks involving risk and cooperation, participants decided whether to take financial risks or to cooperate under possible betrayal. Our results showed that betrayal aversion had a stronger effect on reducing cooperation compared to loss aversion. Furthermore, ERP data demonstrated a temporal pattern consistent with sequential processing: betrayal was encoded early in decision-making, reflected by increased P3 with weaker betrayal aversion, whereas loss aversion manifested later, marked by increased LPP. By dissociating the contributions of betrayal and loss, our finding provides novel insights into the cognitive and neural mechanisms underlying cooperative behavior.

## 1 Introduction

Cooperative behavior refers to activities that benefit another individual or benefit both the person performing the behavior and the recipient (Melis & Semmann, 2010). From collaborating on school projects to managing household responsibilities, humans often engage in cooperative acts that require personal costs (e.g., time, effort, money) for mutual benefit (Tomasello et al., 2005). Such cooperation is central to social life and human interaction, and has shaped human evolution (Nowak, 2006; West et al., 2021). Classic economic games, including the prisoner’s dilemma and the one-shot Trust Game, have provided influential frameworks for studying cooperation, trust, reciprocity, and betrayal (Berg et al., 1995; Fehr & Fischbacher, 2003). These studies show that people often cooperate with others even when betrayal would maximize immediate self-interest, suggesting that cooperative decisions are shaped not only by material incentives but also by social preferences, and expectations about partner trustworthiness (Berg et al., 1995; Fehr & Fischbacher, 2003; Ma et al., 2022). It is widely known that betrayal provokes negative emotions and promotes punishment (Fehr & Gächter, 2000). While much prior research has only focused on “betrayal aversion”, less attention has been paid to the financial loss from unmet expectations during social interactions (Bohnet et al., 2008; Fehr & Fischbacher, 2003). Indeed, mutual cooperation maximizes social welfare, whereas betrayal benefits the trustee but comes at the trustor’s expense in the Trust Game (Berg et al., 1995). However, classic cooperation paradigms such as the prisoner’s dilemma and the Trust Game often involve simultaneous independent choices, repeated interactions, strategic interdependence, and/or sequential reciprocal transfers between partners. Although these features are central to real-world cooperation, they make it difficult to isolate the social meaning of betrayal from the financial loss caused by betrayal. To address this issue, we used a simplified cooperation task structurally matched to a nonsocial risk task to examine whether reduced cooperation is driven primarily by the interpersonal cost of betrayal, the financial cost of potential loss, or both. Crucially, since betrayal and financial loss inherently co-occur during defection, they are difficult to isolate experimentally. Therefore, we use computational modeling to quantitatively disentangle betrayal aversion from loss aversion. Understanding these distinctions is critical for linking computational and neural explanations of cooperative behavior (Krakauer et al., 2017; Lockwood et al., 2020).

Decision uncertainty is typically divided into nonsocial risk (e.g., lotteries) and social risk (e.g., betrayal), where outcomes depend on another individual’s actions (Bohnet & Zeckhauser, 2004). Under nonsocial risk, behavior is often governed by loss aversion, the tendency to weight potential losses more heavily than equivalent gains (Kahneman & Tversky, 1979), which explains why investors typically avoid placing all their eggs in one basket. By contrast, social risk involves not only potential monetary loss but also the possibility of betrayal aversion, defined as the additional aversive response elicited when a loss is caused by another person’s betrayal rather than by chance (Aimone & Houser, 2012). In these studies, participants faced two distinct scenarios: one in which they could keep their money or triple it by giving it to someone else, who might either split it fairly or return only a small portion (less than the original amount), and another involving a lottery-like scenario with identical payoffs, where the return outcome was determined purely by chance. Analyses compared the minimum probability at which people accepted risks between social and nonsocial domains. A consistent finding is that participants are less willing to take risks in the trust game than in the lottery game (Bohnet et al., 2008; Bohnet & Zeckhauser, 2004), an effect termed betrayal aversion (Bohnet & Zeckhauser, 2004). This aversion is modulated by a variety of factors, including individual characteristics such as gender (Croson & Gneezy, 2009; Finkel et al., 2002) and broad personality traits (Komiya & Mifune, 2015; Thielmann & Hilbig, 2015; Yamagishi, 2011), as well as socio-environmental factors like social status (Hong & Bohnet, 2007), and geographic region (Bohnet et al., 2010). Additionally, psychophysiological factors, such as emotional states (Kugler et al., 2012) and oxytocin (Kosfeld et al., 2005), also shape betrayal aversion. Furthermore, the fundamental distinction between social and nonsocial risk extends to their underlying neural mechanisms. Neuroimaging studies reveal that the differences between betrayal costs and loss costs are tracked by neural activity in the amygdala (van Honk et al., 2013) and insula (Aimone et al., 2014). Together, these findings suggest that social and nonsocial risks differ not only in the source of uncertainty and the psychological costs they impose, but also in the neural mechanisms through which these costs are processed (Fehr, 2009a; Lauharatanahirun et al., 2012).

While existing studies have focused on how people choose to cooperate when faced with social and nonsocial risks, they have largely overlooked an essential aspect of betrayal cost: the financial loss inherent in betrayal. Specifically, when investors decide to trust trustees, who may either reciprocate or betray, betrayal simultaneously inflicts both emotional harm and financial loss on the trustor (Aimone & Houser, 2011). This dual nature provides the basis for hypothesizing that betrayal aversion and loss aversion may independently affect cooperation choices. However, prior studies have treated betrayal aversion and loss aversion as separate phenomena by examining them under distinct uncertain conditions. This methodological limitation leaves unresolved whether the observed reluctance to cooperate stems from the psychological impact of betrayal itself or the direct financial loss it causes, even within the same cooperative context. To address this, we used Cumulative Prospect Theory (CPT; Tversky & Kahneman, 1992) to represent the shared monetary prospect. CPT describes risky choice with a value function that weights losses more heavily than equivalent gains (loss aversion, λ) and a probability-weighting function that distorts instructed probabilities. Because the cooperation and risk tasks used identical principals and probabilities, the same CPT value function can represent the financial consequences of accepting versus rejecting the risky choice in both tasks. Betrayal aversion can then be entered as an additional cost on the loss branch of the cooperation task, where the downside is a partner’s defection rather than chance. Comparing a CPT model of monetary value with a model that additionally includes this social cost tests whether reduced cooperation reflects loss aversion, betrayal aversion, or both. Moreover, due to the poor temporal resolution of fMRI, previous studies may have conflated neural activities associated with cognitive processes that occur closely in time but are psychologically distinguishable. Indeed, social decision-making can be decomposed into different stages, including early automatic information processing and late-stage elaborative cognitive evaluation, each with distinct ERP components (Fan & Han, 2008; Wang et al., 2023; Yu et al., 2022).

To address these challenges, we employed the event-related potential (ERPs), which provide excellent temporal resolution to characterize the neural dynamics underlying distinct cognitive processes engaged in social decision-making (Luck, 2014). Cooperative choices are mainly associated with two ERP components: the P3 and the late positive potential (LPP). The P3 is a parietally distributed positive deflection that peaks around 350–500 ms after stimulus onset, which reflects a rapid encoding of motivational significance in prosocial actions (Li et al., 2020; Wang et al., 2017), with larger P3 amplitudes signifying greater prosocial motivation (Chiu Loke et al., 2011; Li et al., 2023; Ma et al., 2011). Using a donation task, Carlson et al. (2016) found that high-versus low-empathy targets increased P3 under intuitive but not reflective decision-making. Directly following the P3, the LPP is a sustained positive deflection over centroparietal areas following the stimulus onset (Hajcak et al., 2009). Unlike the P3, the LPP is thought to reflect the extended, elaborative emotion and motivation processes (Glazer et al., 2018) and might be sensitive to social information evaluation. Crucially, mounting evidence suggests that prosocial actions may stem from fast, automatic, and intuitive processes, while reflective control can reduce them (Rand et al., 2012; Righetti et al., 2013; Zaki & Mitchell, 2013). Together, these findings illustrate the possibility that the P3 may provide an early motivational signal that fosters intuitive prosocial behavior, while LPP may represent a later, more elaborate evaluation that promotes reflective prosocial actions. However, previous ERP research has never considered the joint impact of betrayal and loss aversion on cooperation.

To assess the motivations for cooperation and provide computational and neural accounts of betrayal and loss, we combined computational modeling and ERP with two decision-making tasks: a risk task and a cooperation task. They shared identical probabilities (10%-90%) and principal amounts (¥0.2-¥1.0). This design allowed us to hold probability levels and monetary outcomes constant while varying the source of uncertainty: chance in the risk task versus another person’s betrayal in the cooperation task. They required participants to choose between a risky option, which offered a chance to double the principal at the risk of losing half, and a safe option to retain the initial stake. The primary distinction lies in the social context: in the cooperation task, choosing the safe option constituted non-cooperation, whereas choosing the risky option constituted cooperation, which risked a loss (identical in magnitude to the loss in the risk task) to the partner for potential mutual gain. Participants believed they were interacting with ostensibly real, anonymous human partners, though in reality, all co-player choices were computer-generated to maintain strict experimental control. From the fitted model, the loss aversion coefficient (λ) captured the subjective weight of monetary losses relative to gains, and betrayal aversion coefficient (κ) captured the additional social cost of a partner’s defection. Based on previous studies, we hypothesized that the early P3 reflects intuitive prosocial responses, encoding betrayal aversion, with larger P3 linked to lower betrayal aversion. In contrast, late LPP integrates betrayal and loss aversion after more deliberate evaluation, shown by a more positive LPP as betrayal aversion decreased and loss aversion increased. To provide a comprehensive understanding of cooperation, we also examined participants’ decision-making tendencies across the tasks. We hypothesized that participants would be less willing to take risks facing social risk than nonsocial risk. This study was not preregistered.

## 2 Results

### 2.1 Individuals are also less willing to take risks in the cooperation task than in the risk task

As a manipulation check, we examined participants’ rating data as a function of task type and outcome feedback with a series of repeated-measures analyses of variance (ANOVAs). As shown in Figure 1D, participants rated positive outcomes as more pleasant than negative outcomes, reflected by a significant main effect of outcome feedback, F (1, 41) = 297.01, *p* < 0.001, ηp² = 0.88. As expected, we also found a significant interaction between task type and outcome feedback, F (1, 41) = 16.75, *p* < 0.001, ηp² = 0.29. Post hoc comparisons revealed that the cooperation task elicited greater happiness than the risk task in response to positive feedback (8.29 ± 0.17 vs. 7.76 ± 0.19, *p* = 0.005, Cohen’s d = 0.30). Similarly, the cooperation task elicited greater unhappiness than the risk task with negative feedback (3.12 ± 0.22 vs. 3.83 ± 0.20, *p* = 0.002, Cohen’s d = 0.41). Together, our rating data suggest that the manipulation of the cooperation task was effective.

**Figure 1.**
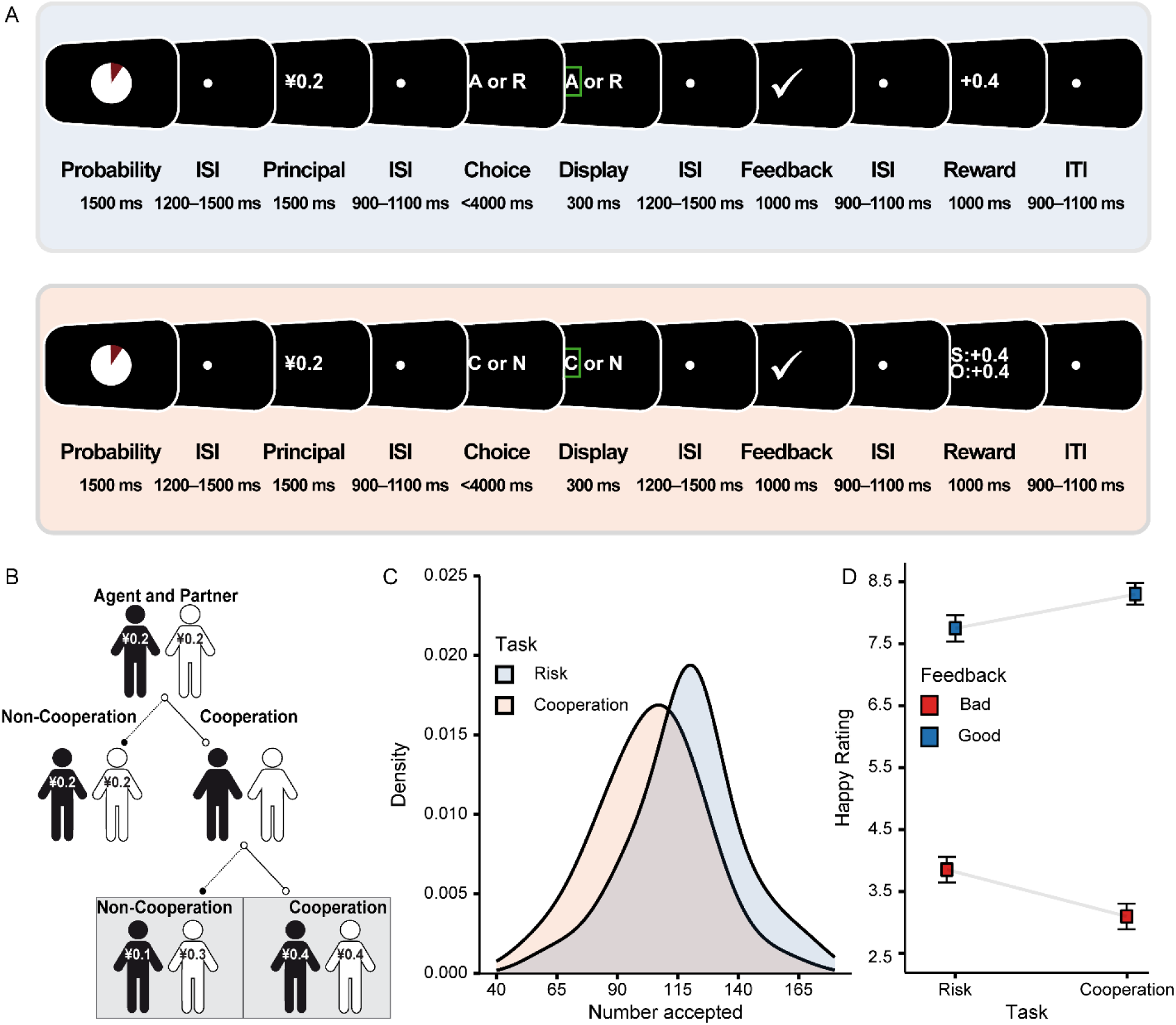
Experimental tasks and rating results. (A) The risk task (top). Participants chose a reject option, keeping their current principal (¥0.2, ¥0.4, ¥0.6, ¥0.8, or ¥1.0), and an “accept” option, where they gambled to double their principal, with a varying probability (10%, 30%, 50%, 70%, 90%) of success, or lose half the principal if they fail. The cooperation task (bottom). Participants as agents chose whether to cooperate with their partner, with the same probabilities and payoffs as the risk task. If successful, both agent and partner doubled their principal; if it failed, the partner took half of the agent’s principal. ISI = interstimulus interval; ITI = intertrial interval. (B) The decision tree depicts an example of a decision to cooperate, and the partner could choose cooperation or betrayal. (C) The distribution of the accepted numbers during the risk task and the cooperation task. (D) Rating data. Participants felt more liking for positive outcomes and more disliking of negative outcomes during the cooperation task than the risk task. Error bars represent the within-subject standard error of the mean.

Because decision time is commonly used as an index of decision difficulty, with longer decision times reflecting greater conflict during value comparison (Evans et al., 2015; Krajbich et al., 2010), we fitted decision-time data using a linear mixed-effects regression with probability, principal, task type, and their interactions as predictors. As illustrated in Figure 2A–C, decision time decreased as probability (*b* = –21.51, *p* < 0.001) and principal magnitude (*b* = – 26.02, *p* < 0.001) increased. Additionally, a significant interaction between probability and principal magnitude was observed (*b* = 12.97, *p* < 0.001). Simple slopes analyses revealed that decision time became shorter as probability level increased at low principal magnitude (*M* –1*SD*: *b* = –34.48, 95% CI = [-43.70, –25.22], *p* < 0.001), but not at high principal magnitude (*M* + 1*SD*: *b* = –8.54, 95% CI = [-17.80, 0.72], *p* = 0.071; direct slope comparison: χ²(1) = 15.08, *p* < 0.001). Decisions were also faster in the cooperation task than the risk task (*b* = –73.89, *p* = 0.002). We also observed significant interactions between probability and task type (*b* = 26.55, *p* < 0.001), as well as principal and task type (*b* = –13.33, *p* = 0.046). Simple slopes analyses revealed that increased probability level decreased the decision time in the risk task (*b* = –34.78, 95% CI = [-44.00, –25.53], *p* < 0.001), but not in the cooperation task (*b* = –8.24, 95% CI = [-17.5, 1.02], *p* = 0.081; direct slope comparison: χ²(1) = 15.80, *p* < 0.001). However, increased principal magnitude decreased the decision time more pronouncedly in the cooperation task (*b* = –32.70, 95% CI = [-41.90, –23.40], *p* < 0.001) than in the risk task (*b* = –19.40, 95% CI = [-28.60, – 10.10], *p* < 0.001; direct slope comparison: χ²(1) = 3.98, *p* = 0.046). Full regression estimates are shown in Supplementary Table S1. We conducted an exploratory expected value (EV)-based reanalysis of decision time. We found a nonlinear relationship where decision times tend to be longer near the objective EV indifference region (Supplementary Table S6).

**Figure 2.**
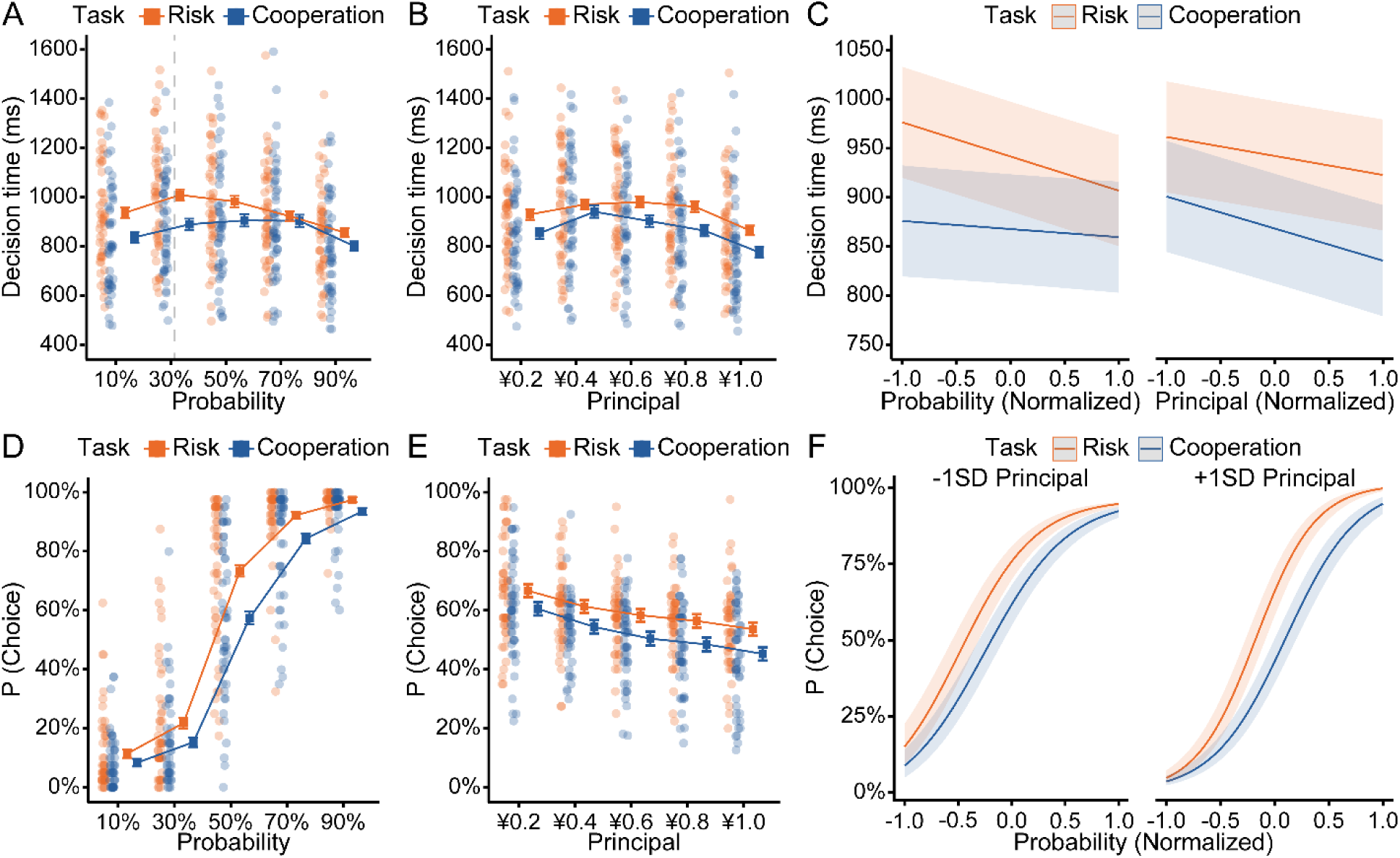
Behavioral results. (A) Participants took shorter to make decisions as the probability level increased in risk task but not in cooperation task. The vertical dashed line marks *p*=1/3, the objective EV-indifference point at which the safe and risky options have equal expected value. (B) Increased principal magnitude decreased the decision time more pronouncedly in cooperation task than in risk task. (C) Fixed effects of probability and principal on the decision time as a function of task type. (D–E) Participants were less willing to take risks in the cooperation task than in the risk task, with the effect being more pronounced at high principal magnitude. (F) Fixed effects of probability on the acceptance rate as a function of task type and principal level (low vs. high). Shaded areas depict the 95% confidence intervals.

Participants’ decision preference was fitted by a mixed-effects logistic regression with probability, principal, task type, and their interactions as predictors. As shown in Figure 2D–E, risk-taking increased as the probability level increased (*b* = 3.00, *p* < 0.001) and principal magnitude decreased (*b* = –0.42, *p* < 0.001). These effects were further qualified by a significant interaction between probability and principal (*b* = 0.31, *p* < 0.001). Simple slopes analyses revealed that participants were more willing to take risks as probability level increased at high principal magnitude (*M* + 1*SD*: *b* = 3.31, 95% CI = [3.00, 3.63], *p* < 0.001) than at low reward magnitude (*M* – 1*SD*: *b* = 2.70, 95% CI = [2.39, 3.01], *p* < 0.001; direct slope comparison: χ²(1) = 93.15, *p* < 0.001). Moreover, they exhibited lower risk-taking in the cooperation task than the risk task (*b* =-0.83, *p* < 0.001), further modified by task type × probability (*b* = –0.44, *p* < 0.001) and task type × principal interaction (*b* = –0.10, *p* = 0.029). Simple slopes analyses revealed that the inhibition effect of the principal on willingness to take risks was more pronounced during the cooperation task (*b* = –0.47, 95% CI = [-0.53, –0.41], *p* < 0.001) compared to the risk task (*b* = – 0.37, 95% CI = [-0.44, –0.30], *p* < 0.001; direct slope comparison: χ²(1) = 4.75, *p* = 0.029). Similarly, the facilitation effect of probability on willingness to take risks was less pronounced during the cooperation task (*b* = 2.78, 95% CI = [2.47, 3.09], *p* < 0.001) compared to the risk task (*b* = 3.23, 95% CI = [2.91, 3.54], *p* < 0.001; direct slope comparison: χ²(1) = 36.63, *p* < 0.001).

The probability × task type interaction was further qualified by a significant three-way interaction among probability, principal, and task type (*b* = –0.20, *p* = 0.002). Post hoc simple slope analyses (Figure 2F) showed that the facilitative effect of probability on risk-taking was attenuated in the cooperation task than the risk task, particularly at high principal magnitude (risk: *b* = 3.63, 95% CI = [3.29, 3.97], *p* < 0.001; cooperation: *b* = 2.99, 95% CI = [2.67, 3.32], *p* < 0.001; direct slope comparison: χ²(1) = 36.37, *p* < 0.001) and to a lesser extent at low principal magnitude (risk: *b* = 2.82, 95% CI = [2.50, 3.14], *p* < 0.001; cooperation: *b* = 2.57, 95% CI = [2.26, 2.89], *p* < 0.001; direct slope comparison: χ²(1) =7.89, *p* = 0.005). Full regression estimates are shown in Supplementary Table S1. Overall, participants were less motivated to take risks in the cooperation task than the risk task, as evidenced by lower acceptance rates and shorter decision times in the cooperation task.

### 2.2 Model-derived betrayal aversion suppresses cooperation willingness to a higher extent than loss aversion

To move beyond observable behavioral effects and distinguish monetary loss sensitivity from the additional social costs associated with partner defection, we applied a family of cumulative prospect theory (CPT)-based models, built in successive steps. Model 0 was a random choice baseline. Model 1 implemented CPT for the shared monetary prospect, with a value function (α_gain, α_loss, λ) and a probability-weighting function (γ) that nonlinearly distorts instructed probabilities. Model 2 added a betrayal-aversion coefficient (κ) on utility function in cooperation task. This ladder tests whether choices were accounted for by CPT monetary value, or by an additional social cost of partner defection (see Method for more details). Model recovery showed that these models produced distinguishable choice patterns, with recovery accuracy ranging from 80% to 100% (Supplementary Figure S2). Model comparison decisively favored Model 2 (AIC = 240.7, BIC = 264.7, CVIC = 243.4; versus 250.4, 270.4, 253.0 for Model 1 and 533.0, 536.9, 533.2 for Model 0; Figure 3A; Table S2). Parameter recovery analysis showed the parameters of M2 were well identifiable (α_gain = 80%, α_loss = 93%, γ = 89%, λ = 89%, β = 84%, κ = 81%; Figure 3B; details of model comparison and parameter recovery see Supplementary Table S2 and S3). Parameter-recovery plots for Models 0–2 are shown in Supplementary Figures S1. In a separate model-based simulation, participant-level parameter estimates from Model 2 reproduced the observed effects of probability and principal magnitude on acceptance in both tasks (Supplementary Figure S3).

**Figure 3.**
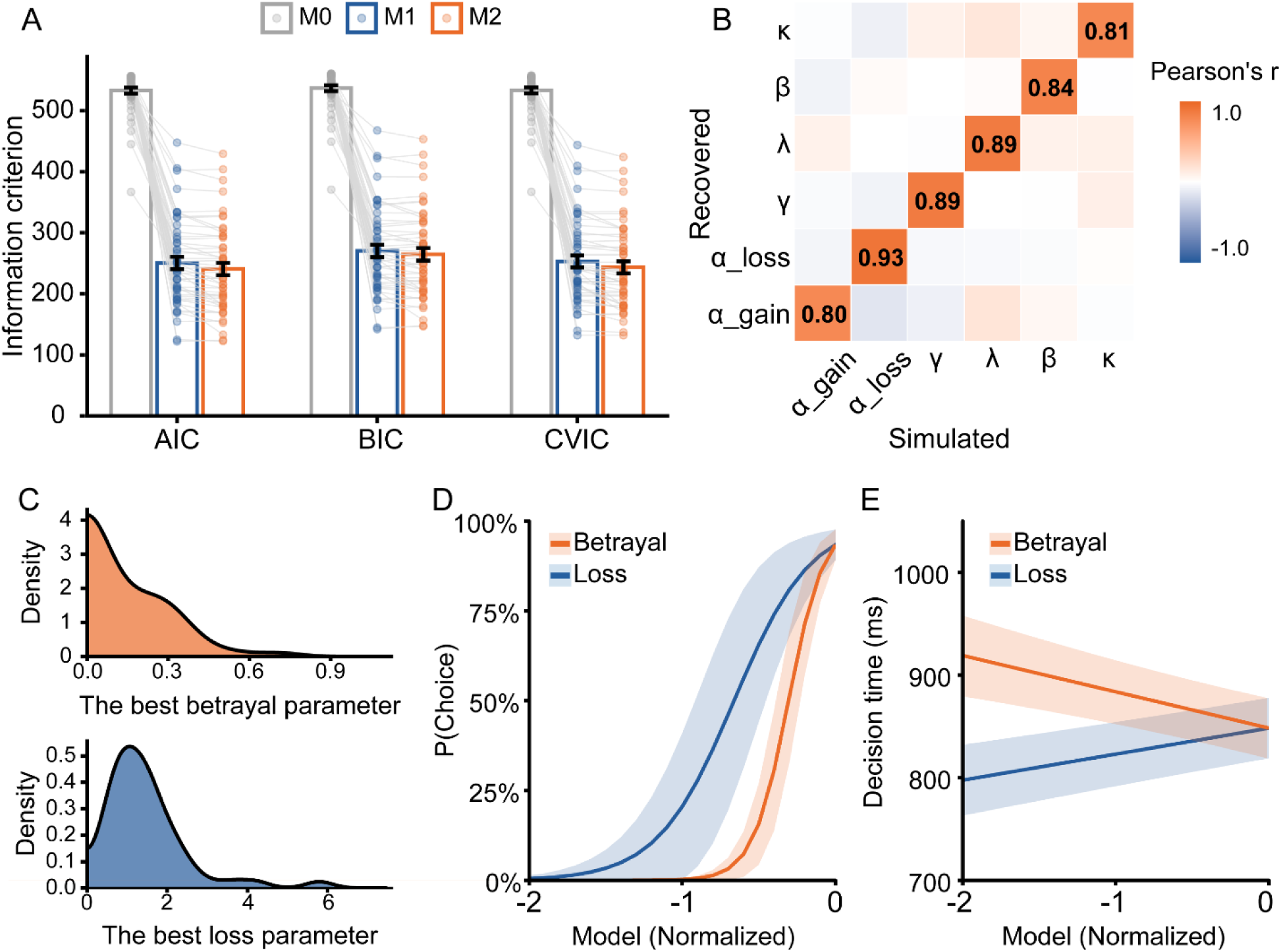
Computational results. (A) Mean AIC, BIC, and CVIC values (bars; lower values indicate better fit) from the comparison of the computational models. Points represent individual participants. Model 2 (incorporating CPT, betrayal aversion) was selected as the winning model. (B) Parameter recovery of Model 2: gain sensitivity (α_gain), loss sensitivity (α_loss), probability sensitivity (γ), loss aversion (λ), noise (β), and betrayal aversion (κ). The correlation matrix shows the Pearson correlation between the actual recovered and simulated data from the winning model. The high correlations (all *r* > 0.80) indicate excellent parameter recovery. (C) The distribution of the best betrayal (top) and loss (bottom) parameters for the winning model, which maximizes log-likelihood during fitting. (D–E) The willingness to cooperate became lower as loss aversion and betrayal aversion increased (D), and decision time became shorter as loss aversion increased and betrayal aversion decreased (E). Betrayal aversion significantly predicts both acceptance rate (D) and decision time (E). Shaded areas depict the ± 1 standard errors. Loss represents the subjective value for the best-fitting financial loss aversion (−λ ∗ (0.5 ∗ V)^αloss^ ∗ *P_Weig_*_ℎ*t*_) and betrayal is the subjective value of betrayal aversion κ ∗ *P_Weig_*_ℎ*t*_). \*\*\**p* < 0.001, \*\**p* < 0.01.

To examine how betrayal aversion and loss aversion influence cooperation behavior, we fitted decision time using a linear mixed-effects regression and decision preference using a mixed-effects logistic regression with a binomial link. As shown in Figure 3D, both loss aversion and betrayal aversion significantly predicted cooperation, such that participants cooperated less as loss aversion (*b* = 4.00, *p* < 0.001) and betrayal aversion (*b* = 8.66, *p* < 0.001) increased. Moreover, we employed β coefficient tests to compare the predictive power of loss and betrayal aversion on cooperation willingness, revealing that betrayal aversion had a stronger impact on cooperation than loss aversion (*z* = –3.36, *p* < 0.001). As shown in Figure 3E, both aversions also robustly predicted decision times, with decision time becoming shorter as loss aversion increased (*b* = 25.37, *p* < 0.001) and betrayal aversion decreased (*b* = –35.35, *p* = 0.023), again with betrayal aversion showing greater predictive capacity (*z* = 3.53, *p* < 0.001). Full regression estimates are provided in Supplementary Table S4. Together, our computational modeling indicates that betrayal aversion has a stronger suppressive effect on cooperative choices than loss aversion, reflected in lower cooperation willingness and longer decision times.

### 2.3 Betrayal-related information was associated with earlier decision processing, reflected in the P3, whereas loss information was incorporated later, indexed by the LPP

The P3 and LPP components were observed as a relative positivity over centroparietal areas (Figure 4). Betrayal– and loss-aversion parametric were derived from the winning model (Model 2). To investigate their effects on neural dynamics underlying cooperation, we fitted the amplitude of the P3 and LPP using a linear mixed-effect regression with z-scored betrayal and loss parametric as predictors. As shown in Figure 4, the P3 became less positive as betrayal aversion increased (*b* = 0.45, *p* = 0.015), but was unaffected by loss aversion (*b* = –0.07, *p* = 0.57). A β-coefficient comparison confirmed that the effect of betrayal aversion was more significant (*z* = –2.37, *p* = 0.009), indicating a betrayal inhibition effect. For the LPP, positivity decreased as betrayal aversion increased (*b* = 0.68, *p* < 0.001) and loss aversion decreased (*b* = −0.33, *p* = 0.009). The β-coefficient comparison further showed that betrayal aversion had a stronger predictive effect than loss aversion (*z* = –4.57, *p* < 0.001). Full regression estimates are shown in Supplementary Table S5.

**Figure 4.**
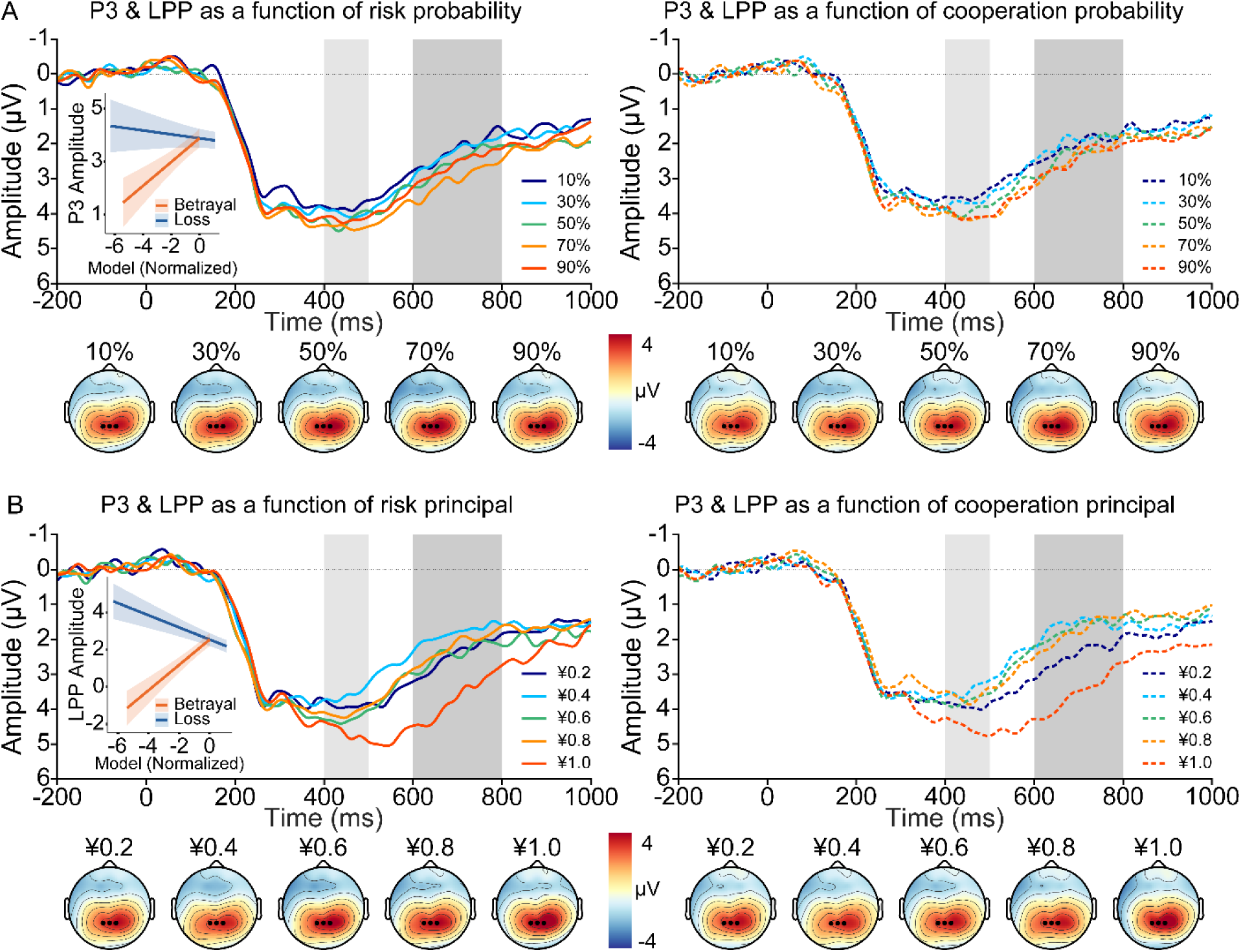
Grand-average ERP waveforms and topographic maps of the P3 and LPP as a function of task type (risk task vs. cooperation task) separately for probability. (A) and principal (B) trials. Gray shaded bars represent time windows used for quantification. The insets show the fixed effects of betrayal and loss aversion on the P3 (A) and LPP (B). The betrayal-inhibition effect emerged earlier and persisted later, reflected in both P3– and LPP-range activity, whereas the loss-inhibition effect appeared only later, reflected in LPP-range activity. Shaded areas depict the 95% confidence intervals.

Together, our ERP data are consistent with a temporal account in which betrayal-related and loss information were processed serially, such that defection-related information was associated with the earlier P3-range activity, whereas loss information was associated with the later LPP-range activity.

## 3 Discussion

Betrayal suppresses cooperation and trust, yet the associated financial losses it incurs are often overlooked, complicating the reasons behind cooperative behavior. Here, we explore how betrayal and its financial consequences influence cooperation alongside their underlying neural mechanisms, addressing a gap in previous research. We found that betrayal exerts a significantly stronger suppression on cooperation than loss. Moreover, neural processing unfolded sequentially: betrayal modulated early-stage automatic processing indexed by the P3, while loss appeared in later evaluative stages captured by the LPP.

In our study, participants reported a stronger preference for positive feedback and greater aversion to negative feedback during the cooperation task, indicating stronger affective responses to outcomes in the cooperation task than in the risk task. They were also less willing to take risks as the cooperation risk increased, especially when the principal magnitude was high, compared to the nonsocial risk. Moreover, in the cooperation task, decision times were shorter and driven more by principal amounts rather than probability levels, indicating that participants prioritized potential gains and losses and adopted a generally cautious approach, regardless of the cooperative probability. These findings are largely consistent with previous studies, which compared individuals’ willingness to take risks when the outcome depends on another player’s trustworthiness, versus situations where the outcome is determined by a chance with identical odds and payoffs, demonstrating a stronger avoidance motivation in social contexts (Bohnet et al., 2008; Bohnet et al., 2010; Bohnet & Zeckhauser, 2004). However, these studies overlook that betrayal simultaneously incurs emotional and financial losses.

By employing computational modeling to separately examine how betrayal and its associated loss parameters influence cooperation, our most important finding is a greater substantial inhibitory effect of betrayal on cooperation than loss. Participants exhibited lower cooperation rates as both loss aversion and betrayal aversion increased and longer decision times as loss aversion decreased and betrayal aversion increased. Notably, betrayal aversion had a stronger impact on choice or decision time. Several theoretical frameworks can help interpret this effect. First, individuals care about others’ payoffs in cooperative situations (Fehr, 2009b). In our study, when partners betrayed by taking half of the investor’s money, participants are less likely to accept the social risk compared to the nonsocial risks. Second, people care not only about outcomes but also the intentions behind them (Charness & Rabin, 2002; Rabin, 1993). Trust builds honor and integrity, promoting cooperation, while betrayal erodes these qualities, discouraging future cooperation. Third, betrayal often inflicts emotional harm, such as damaged trust, which can outweigh material loss. Aimone et al. (2014) found stronger activation in the anterior insular cortex when playing with a human versus a computer, indicating heightened negative emotions (Kuhnen & Knutson, 2005; Wu et al., 2012), supporting betrayal aversion as a desire to avoid negative emotions. Finally, people perceive risks from others’ actions as less controllable than natural risks (Slovic, 1987). For example, shareholders would prefer a 1% chance of losing half their value due to a natural disaster than a slightly smaller chance from betrayal by corporate executives. Here, betrayal aversion (κ) captures the additional cost of a partner’s defection beyond monetary loss. This cost may reflect unfairness or perceived intent, which our model does not distinguish.

Interestingly, we observed distinct neural mechanisms for betrayal and loss. Specifically, the early P3 amplitude decreased with increasing betrayal aversion, while no significant change was observed in response to loss aversion. In contrast, LPP amplitude became less positive with stronger betrayal aversion but more positive with stronger loss aversion. These findings suggest that betrayal and loss information are processed sequentially, with betrayal being encoded during the P3 period, followed by further integration of loss information during the LPP period. Cooperation is often considered an intuitive behavior (Gao et al., 2020; Koch et al., 2020; Levine et al., 2018; Rand et al., 2012; Shank et al., 2019), which is indexed by P3, a neural signal that rapidly encodes prosocial motivation (Li et al., 2020; Wang et al., 2017). Betrayal, a key factor inhibiting prosocial behavior (Fehr & Gächter, 2000), is therefore prioritized for encoding at the early stages of cooperation decision-making. According to a dual-process framework, intuition is fast, automatic, and effortless, while reflection is often slow, deliberate, and characterized by the rejection of emotional influence (Chaiken & Trope, 1999; Frederick, 2005; Kahneman, 2011; Plessner et al., 2008; Sloman, 1996; Stanovich & West, 1998). Hence, during the later phase of decision-making, participants further engage in a trade-off between costs and benefits associated with specific actions. This process is indexed by the LPP, a neural marker reflecting sustained and elaborative information processing (Glazer et al., 2018). In this stage, loss-related information is integrated into the decision-making process. Thus, the P3-range activity should be interpreted as an early ERP correlate of defection-related social cost during cooperation decisions, rather than as a specific neural marker of betrayal aversion.

A limitation of this study is the ecological validity of the cooperative scenario. Participants were informed that they would cooperate with others, but some (N=6) expressed doubts about the authenticity of this interaction in a post-experiment check. Although this did not affect the main results, future studies could investigate neural synchronization during multi-player cooperation to better understand how co-players synchronize their neural activity in such contexts (Jiang et al., 2015; Yang et al., 2020; Zhang et al., 2023). Additionally, future studies could provide evidence to distinguish the aversion to being personally betrayed and witnessing another’s betrayal. This approach will help determine whether a betrayal has a stronger inhibitory effect and is processed with priority from a third-party perspective, where the outcome is unrelated to oneself. Finally, we only recorded general happiness ratings, which cannot directly quantify specific emotional costs like betrayal. Additionally, we did not assess baseline individual traits. Future studies should incorporate trial-by-trial ratings of perceived betrayal and trait-level measures, such as Social Value Orientation (SVO), to clarify how specific emotions and individual differences jointly modulate decision-making. More broadly, because this study was not preregistered, the findings should be interpreted with appropriate caution and further validated in preregistered studies.

In conclusion, this study demonstrates that betrayal has a stronger suppressive effect on cooperation compared to the associated cost of losses. Additionally, this study provides preliminary evidence that the neural dynamics involved in these two forms of aversion occur sequentially, with betrayal aversion influencing early-stage decision-making and loss aversion affecting later-stage evaluations. These findings enhance our understanding of how emotional and financial considerations influence cooperative behavior and provide important insights into the neural mechanisms that underlie cooperative decision-making.

## 4 Materials and methods

All data and code used for this study are available on GitHub at https://github.com/jbgor/2026_eLife_Betray_aversion

### 4.1 Participants

Fifty right-handed university students were recruited via local advertisements for this study. One participant was excluded from data analysis due to a misunderstanding of instructions. The final sample thus consisted of 49 participants (24 females; *M* = 22.00 years, standard deviation [*SD*] = 2.03). All participants had normal or corrected-to-normal vision as determined by self-report and no psychiatric or neurological disorders. Each received a bonus of ¥70–¥80 based on their task performance. This study was approved by the Institutional Review Board of Sun Yat-sen University.

### 4.2 Procedure

Upon arrival at the lab, participants completed a probabilistic risk task and a single-shot cooperation task while their EEG was recorded. After the EEG tasks, participants rated their perceived happiness on a 9-point Likert scale (ranging from 1 = not at all to 9 = very much), based on the outcomes (gain or loss) associated with each decision.

#### The risk task

This task was designed to assess neural correlates of loss aversion. On each trial (Figure 1A), participants first viewed a probability pie chart for 1500 ms, followed by a jittered interval (1200–1500 ms). The pie chart illustrated probability levels (10%, 30%, 50%, 70%, or 90%, corresponding to 1, 3, 5, 7, or 9 segments of the chart), indicating the likelihood of doubling the principal versus losing half of it. Afterward, a number appeared for 1500 ms, showing the principal amount (¥0.2, ¥0.4, ¥0.6, ¥0.8, or ¥1.0, corresponding to 1–5 principal levels). The five probability levels were fully crossed with the five principal levels, resulting in 25 unique combinations. Following another jittered interval (900–1100 ms), participants entered the choice phase, choosing between the “reject” option (R), which allowed them to keep the current principal and proceed to the next trial, and the “accept” option (A), which offered a chance to double the principal with the given probability or lose half in case of failure. Participants had up to 4000 ms to decide using their left or right index finger. The chosen option was highlighted with a green border for 300 ms. Failure to respond within the time limit resulted in ¥0 and a 1000-ms warning. After another jittered interval (1200–1500 ms), an outcome feedback stimulus was presented for 1000 ms: a tick denoted gain, a cross denoted loss, followed by a jittered interval of 900–1100 ms. Subsequently, a reward feedback stimulus was shown for 1000 ms, indicating the final payoff. Each trial ended with an interval varying between 900 and 1100 ms.

#### The cooperation task

This task was designed to measure participants’ neural responses to both loss aversion and betrayal aversion, which employed a similar structure but incorporated social decision-making elements. One key distinction was that each trial involved an anonymous co-player, who was distinct from the ones in any other trials and would not reappear in any subsequent trial. Another key difference was the presence of a probability cue reflecting the likelihood that the co-player would cooperate. Thus, lower cooperation probability indicated higher anticipated risk of betrayal, but did not itself constitute betrayal. Participants were informed that the probabilities were derived from prior behavioral data, but in reality, they were randomly generated. Critically, unlike the risk task, where outcomes depended solely on chance, the cooperation task introduced interpersonal contingencies. During the choice phase, participants decided whether to cooperate (C) or not (N). Feedback then revealed the co-player’s decision (a tick for cooperation, a cross for betrayal), followed by a reward screen showing the monetary outcome for both players. Mutual cooperation resulted in the principal being doubled and split equally. If the participant cooperated but the co-player defected, this outcome constituted a betrayal. As a result, half of the participant’s principal was transferred to the co-player. If the participant chose not to cooperate, the principal remained unchanged regardless of the co-player’s choice. The risk task and cooperation decision-making task each consisted of 200 trials, divided into eight blocks of 25 trials, with a self-determined break between blocks. The task order was counterbalanced across participants. To maintain attention, each task included 10 catch trials in which participants were required to confirm the probability and principal levels for the current trial. Before the experiment, participants completed 7 practice trials for each task for familiarization. After the task, participants were asked to report whether they believed they were playing with the third-party players. Three subjects had an accuracy rate of less than 60% on catch trials, and six subjects expressed doubts about the cooperation manipulation, one of these participants had already been excluded from all analyses because of misunderstanding the task instructions. Sensitivity analyses excluding these participants (N = 41) supported the main cooperation-choice and ERP findings, although some decision-time effects no longer reached statistical significance (see Supplementary Tables S7–S12).

### 4.3 EEG recording and processing

EEG data were recorded using 64 Ag/AgCl channels placed on an elastic cap based on the international 10–20 system. Two additional channels were positioned on the left and right mastoids. The reference electrode was located at Cz. Horizontal and vertical electrooculograms were recorded from two pairs of channels over the external canthi of each eye and the left suborbital and supraorbital ridges, respectively. EEG signals were amplified using a Neuroscan SynAmps^2^ amplifier with a low-pass filter of 100 Hz in DC acquisition mode and digitized at a rate of 1000 samples per second. Channel impedances were maintained below 10 KΩ.

The EEG data were analyzed using EEGLAB v2021.0 (Delorme & Makeig, 2004) and ERPLAB v8.10 (Lopez-Calderon & Luck, 2014) toolboxes in MATLAB 2020b (MathWorks, US). The signals were rereferenced to the average of the left and right mastoids and filtered with a bandpass of 0.1–35 Hz using a zero phase-shift Butterworth filter (12 dB/octave roll-off). Channels with poor quality or excessive noise were interpolated using the spherical interpolation algorithm, and portions of EEG containing extreme voltage offsets or break periods were removed. Ocular artifacts were removed using an infomax independent component analysis on continuous EEG with the help of the ICLabel algorithm (Pion-Tonachini et al., 2019). Epochs were then extracted from –200 to 1000 ms relative to principal stimulus onset, which was chosen because it was the first point at which both probability and monetary magnitude were available, with the prestimulus average activity as the baseline. An automatic artifact detection algorithm was applied to remove epochs with a voltage difference exceeding 50 μV between sample points or 200 μV within a trial, a maximum voltage difference less than 0.5 μV within 100-ms intervals, or a slow voltage drift with a slope greater than ± 100 μV. On average, 96.30% of trials were retained for statistical analysis. Measurement parameters were determined by the grand-averaged ERP waveforms and topographic maps collapsed across all conditions (Luck & Gaspelin, 2017). Specifically, the single-trial P3 amplitude and LPP amplitude were measured as the mean activity from 400 to 500 ms and 600 to 800 ms, respectively, after the onset of the principal cues over centroparietal areas (P1, Pz, P2). These single-trial data were exported into R v4.2.2 for statistical analyses.

### 4.4 Data analysis

#### Statistical analysis

Unless otherwise specified, all behavioral, computational-modeling, and ERP analyses included 49 participants. The post-experimental rating analysis included 42 participants because rating data from seven participants were unavailable due to recording problems. Our key statistical analyses were based on mixed-effects regression models with random intercepts and slopes (unstructured covariance matrix), implemented in the *lme4* package v1.1.31 (Bates, Maechler, et al., 2015). For decision time and choice data, we separately used a linear mixed-effects regression model and a mixed-effects logistic regression model with a binomial link function. Both models included probability, principal, task type, and their interactions as predictors. For all models, categorical regressors (task type: –0.5 for risk and +0.5 for cooperation) were contrast-coded, and continuous regressors (probability and principal) were z-scored before inclusion in the regression models. We determined random effects for each model using singular value decomposition to report the maximal possible random effects structure (Barr et al., 2013; Bates, Kliegl, et al., 2015). Follow-up pairwise comparisons of significant interactions were tested on estimated marginal means using the *emmeans* package v1.8.3 (Lenth, 2022). We then evaluated simple-effect differences through linear contrasts via the *linearHypothesis* function in the *car* package v3.1.3 (Fox & Weisberg, 2019). To examine the relationship between behavioral indices of loss aversion (λ) and betrayal aversion (κ) and neural responses to principal cues, both λ and κ were entered as fixed effects in separate P3 and LPP models. All model predictors were scaled (e.g., λ/sqrt(sum(λ^2)/length(λ)-1)) without mean-centering to ensure comparability of β coefficients. We analyzed post-experimental rating data using a repeated-measures analysis of variance (ANOVA), with feedback (good, bad) and task type (risk, cooperation) as within-subjects factors. We excluded trials with no responses (0.15%) in the risk and cooperation task from statistical analyses.

#### Computational modeling

To quantitatively dissociate the contributions of loss aversion and betrayal aversion to cooperative decision making, we employed a computational framework grounded in Cumulative Prospect Theory (CPT; Tversky & Kahneman, 1992). For the participant’s own monetary outcomes, the safe option had an objective EV of X, whereas the risky option had an objective EV of *p*(2X) + (1-*p*) (0.5X). Thus, the two options were equal in objective EV when *p* = 1/3, regardless of principal magnitude. Unlike objective EV, which is determined solely by the instructed probabilities and monetary outcomes, the subjective value estimated by our computational model additionally incorporates probability weighting, gain and loss sensitivity, loss/betrayal aversion, and choice stochasticity. This approach models risky decisions through two core transformations: a value function that asymmetrically weights losses versus gains, and a probability weighting function that nonlinearly distorts objective probability. A family of three models (M0-M2) was systematically constructed to incorporate socially specific factors and test hypotheses about how financial losses and betrayal aversion independently affect cooperation choices. The simplest model (M0) served as a random choice baseline. Model 1 was adapted from the cumulative prospect theory, and defined subjective value (SV) as the net expected utility of choosing investment or cooperation. Specifically, for a given option with principal amount *X* and success probability *p*, the SV was computed as

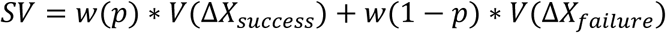

Where the net utility of success Δ*X_success_* = *X* and net utility of *failure* Δ*X_failure_* = −0.5 ∗ *X* as the setting of the experiment. The subjective utility function *V*(*x*) was defined as

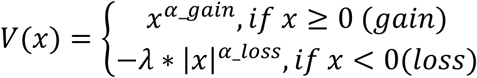

Here, the parameter *α_gain_* and *α_loss_* (0 < *α* < 1) represents the diminishing sensitivity to increasing magnitudes of gains and losses, respectively; *λ* quantifies financial loss aversion, *λ* ≥ 1 indicates that the reduced utility of losses is greater than the utility of gains at the same magnitude.

Besides, the subjective probability weighting function *w*(*p*) was defined as:

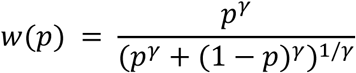

Parameter *γ* (*γ* < 1) governing the curvature and sensitivity of probability for the outcomes. For the simplicity of the model, we estimated a single *γ* parameter across gains and losses. Lower values on *γ* indicate greater curvature and lower sensitivity to probabilities.

Building on M1, Model 2 introduced betrayal aversion as an additional social cost term independent of loss aversion. The SV of M2 was computed as

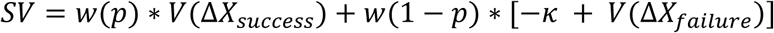

Here, *κ* (*κ* > 0) captures betrayal aversion independent of the monetary loss magnitude, reflecting the hypothesis that social betrayal carries intrinsic disutility beyond monetary consequences.

The derived SV of the option was transformed into choice probability through a softmax function incorporating a stochasticity parameter *β*:

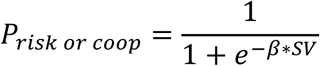

Where *β* (*β* ≥ 0) represents the slope of the logistic function and reflects the sensitivity to SV differences between options. The lower *β* is, the greater the stochasticity of the choices; with *β* = 0, choices are random (Collins & Shenhav, 2022).

For participant *i* and trial *t*, the choice (with *y* = 1 for accept and *y* = 0 for reject) was treated as an independent Bernoulli outcome and the log-likelihood was *ll_it_* = *y_it_* ∗ *logP_it_* + (1 − *y_it_*) ∗ *log*(1 − *P_it_*). The log-likelihood of a model for each participant was *LL_i_* = ∑*_t_ ll_t_*. Parameters of each model were estimated separately for each participant by maximizing likelihood with L-BFGS-B algorithm, and using 20 random initialization points to avoid local minima.

Model comparison used three information criteria computed from each participant’s maximum likelihood and then averaged across participants. With k free parameters and n trials, Akaike Information Criterion (AIC) was −2 ∗ *ll* + 2*k*, Bayesian Information Criterion (BIC) was −2 ∗ *ll* + *k* ∗ ln *n*. We additionally computed a 10-fold cross-validated information criterion (CVIC; Kohavi, 1995; Wilson & Collins, 2019). For each participant, trials were randomly partitioned into 10 folds. On each fold, parameters were re-estimated from the remaining nine folds and the likelihood was evaluated on the held-out trials. The participant level CVIC was −2 ∗ ∑*_folds_ ll*_ℎ*eld*_*out*_. The model with the lowest mean AIC, BIC, and CVIC was selected. The details of information criteria are reported in Supplementary Table S2.

After parameter estimation, the fitted parameters were substituted into Model 2 to compute trial-level values of loss aversion and betrayal aversion,

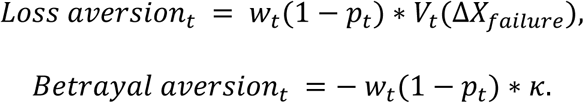

These trial-level values for each participant were then scaled to *zloss* and *zbetrayal,* with more negative utility values indicating stronger loss aversion and betrayal aversion, respectively.

Moreover, the parameters’ identifiability was validated through recovery analyses. For each model, 500 plausible parameter sets were randomly sampled from uniform distributions bounded by empirically informed ranges. Using these ground-truth parameters, we simulated trial-wise choice data (500 trials/set) under experimental conditions matching the original task design. The model was subsequently refitted to the simulated datasets using 20 randomized initialization points per recovery to mitigate local minimum convergence. Parameter recovery similarity was quantified through Pearson correlations between true and estimated parameter values across all simulations.

To test whether Models generate distinguishable choice patterns, we conducted a group-level model recovery. For each true model, 50 synthetic group data sets were generated. In each data set, choices for all 49 participants were simulated from that model, using each participant’s observed trial structure (instructed probability, principal). True parameter vectors were drawn independently for each simulated participant from uniform distributions over the same bounds used in estimation. Simulated choices were independent Bernoulli draws from the model’s trial-wise choice probabilities. Each synthetic data set was then fitted with candidate Model 0, 1, and 2, using the same estimation procedure as the empirical fits (L-BFGS-B; 20 random starting points per participant). For every candidate model, a 10-fold CVIC was computed for each simulated participant and summed across the 49 participants. The candidate with the lowest summed CVIC was selected. Recovery accuracy was the proportion of the 50 data sets in which the selected model matched the true model (Figure S2).

## Acknowledgements

This work was supported by the Natural Science Foundation of Guangdong Province, China (Grant No. 2024A1515010826) and National Natural Science Foundation of China (Grant No. 32571254, 32171073)

## Supplementary Materials

**Table S1.**
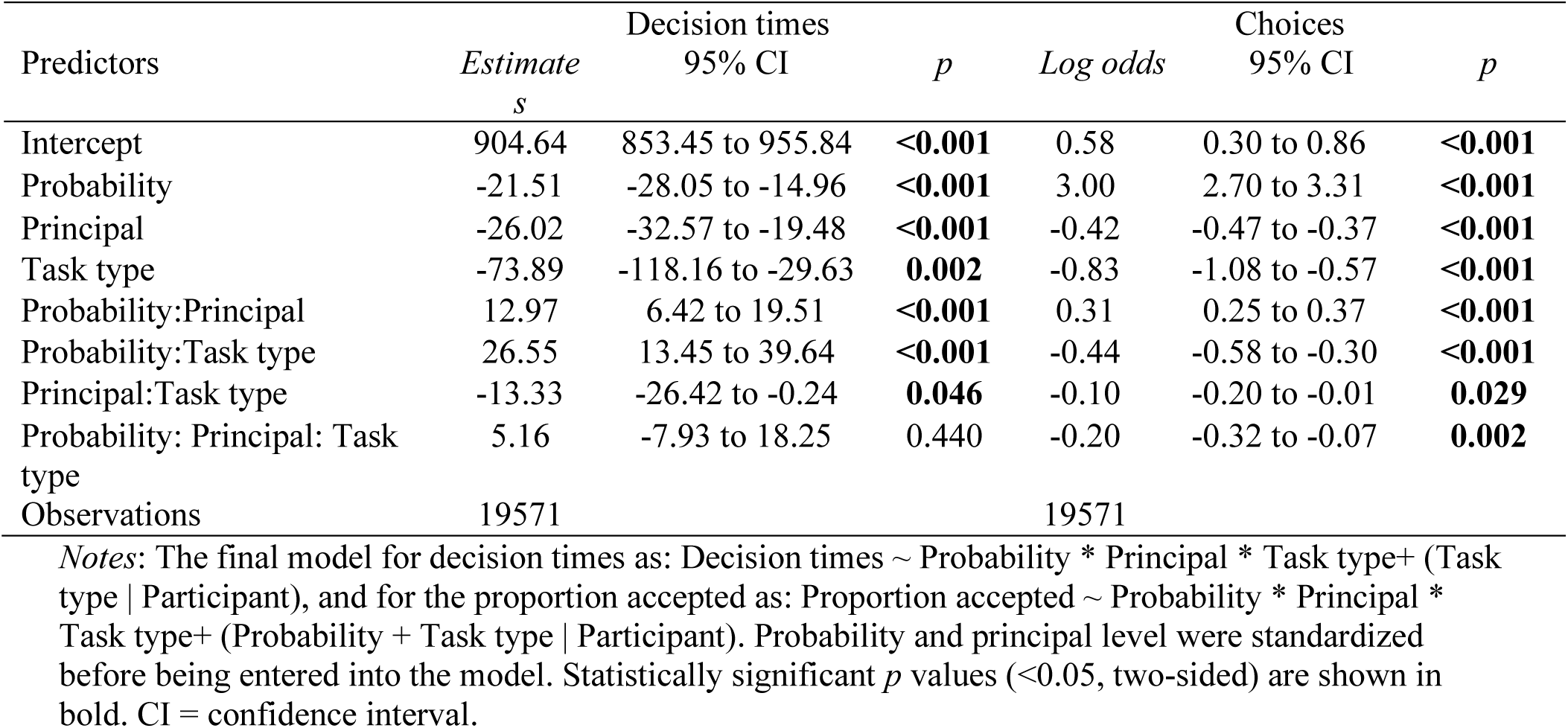
Results of linear mixed-effects models predicting decision times (left) and choices (right) as a function of probability, principal, and task type. (*N*=49)

| Predictors | Decision times |  |  | Choices |  |  |
|---|---|---|---|---|---|---|
|  | *Estimates* | 95% CI | *p* | *Log odds* | 95% CI | *p* |
| Intercept | 904.64 | 853.45 to 955.84 | **<0.001** | 0.58 | 0.30 to 0.86 | **<0.001** |
| Probability | -21.51 | -28.05 to -14.96 | **<0.001** | 3.00 | 2.70 to 3.31 | **<0.001** |
| Principal | -26.02 | -32.57 to -19.48 | **<0.001** | -0.42 | -0.47 to -0.37 | **<0.001** |
| Task type | -73.89 | -118.16 to -29.63 | **0.002** | -0.83 | -1.08 to -0.57 | **<0.001** |
| Probability:Principal | 12.97 | 6.42 to 19.51 | **<0.001** | 0.31 | 0.25 to 0.37 | **<0.001** |
| Probability:Task type | 26.55 | 13.45 to 39.64 | **<0.001** | -0.44 | -0.58 to -0.30 | **<0.001** |
| Principal:Task type | -13.33 | -26.42 to -0.24 | **0.046** | -0.10 | -0.20 to -0.01 | **0.029** |
| Probability: Principal: Task type | 5.16 | -7.93 to 18.25 | 0.440 | -0.20 | -0.32 to -0.07 | **0.002** |
| Observations | 19571 |  |  | 19571 |  |  |

*Notes*: The final model for decision times as: Decision times ~ Probability \* Principal \* Task type+ (Task type | Participant), and for the proportion accepted as: Proportion accepted ~ Probability \* Principal \* Task type+ (Probability + Task type | Participant). Probability and principal level were standardized before being entered into the model. Statistically significant *p* values (<0.05, two-sided) are shown in bold. CI = confidence interval.

**Table S2.** Model comparison (*N*=49)

| Model | Model description | *k* | Mean *AIC* | Mean *BIC* | Mean *CVIC* |
|---|---|---|---|---|---|
| Model 0 | Random model | 1 | 533.0 | 536.9 | 533.2 |
| Model 1 | Cumulative Prospect Theory (CPT) | 5 | 250.4 | 270.4 | 253.0 |
| Model 2 | CPT + betrayal aversion | 6 | 240.7 | 264.7 | 243.4 |

Note: *k* = the number of free parameters; LL = log likelihood; AIC = Akaike information criterion; BIC = Bayesian information criterion; CVIC = 10-fold cross-validated information criterion. Mean AIC, mean BIC, and mean CVIC are averages across participants. Lower values indicate better fit.

**Table S3.**
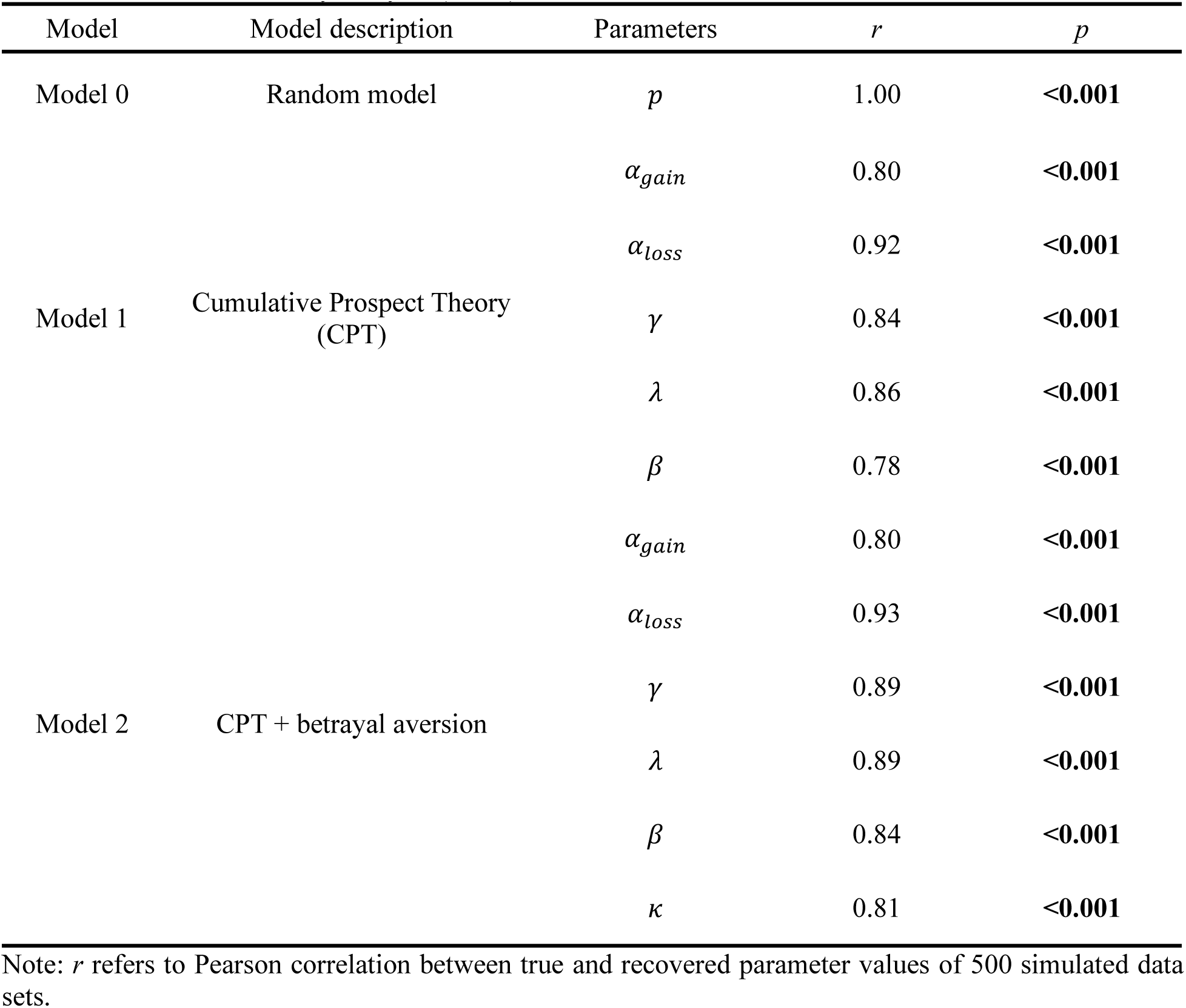
Parameter recovery analysis (*N*=49)

| Model | Model description | Parameters | *r* | *p* |
|---|---|---|---|---|
| Model 0 | Random model | *p* | 1.00 | **<0.001** |
| Model 1 | Cumulative Prospect Theory (CPT) | *α<sub>gain</sub>* | 0.80 | **<0.001** |
|  |  | *α<sub>loss</sub>* | 0.92 | **<0.001** |
|  |  | *γ* | 0.84 | **<0.001** |
|  |  | *λ* | 0.86 | **<0.001** |
|  |  | *β* | 0.78 | **<0.001** |
| Model 2 | CPT + betrayal aversion | *α<sub>gain</sub>* | 0.80 | **<0.001** |
|  |  | *α<sub>loss</sub>* | 0.93 | **<0.001** |
|  |  | *γ* | 0.89 | **<0.001** |
|  |  | *λ* | 0.89 | **<0.001** |
|  |  | *β* | 0.84 | **<0.001** |
|  |  | *κ* | 0.81 | **<0.001** |

Note: *r* refers to Pearson correlation between true and recovered parameter values of 500 simulated data sets.

**Table S4.**
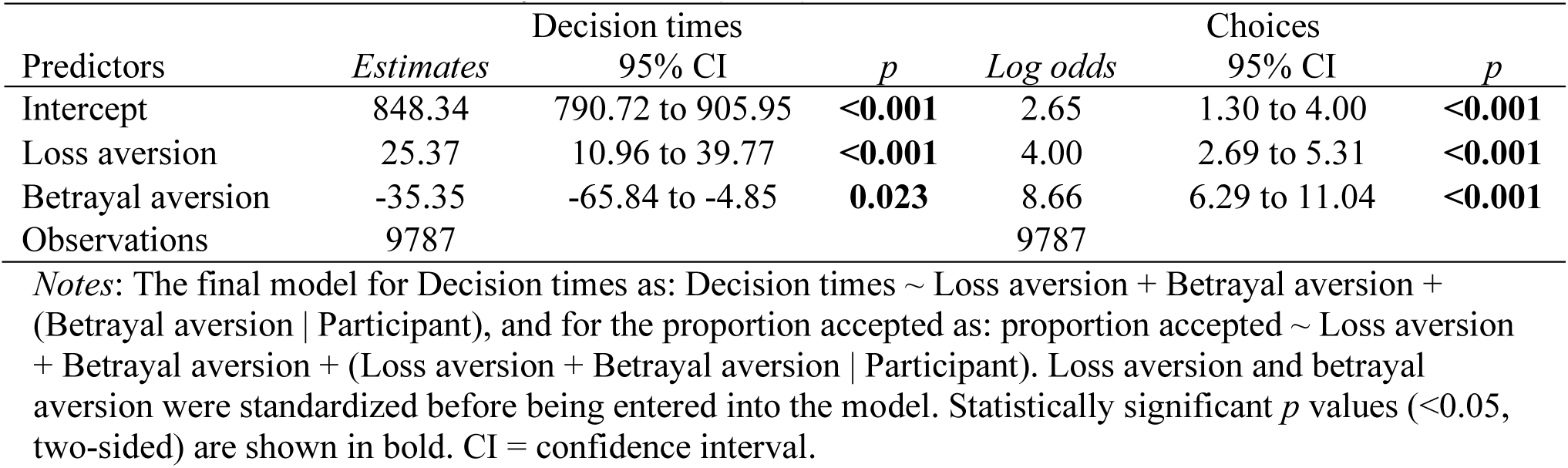
Results of linear mixed-effects models predicting decision times (left) and choices (right) as a function of loss aversion and betrayal aversion. (*N*=49)

| Predictors | Decision times |  |  | Choices |  |  |
|---|---|---|---|---|---|---|
|  | *Estimates* | 95% CI | *p* | *Log odds* | 95% CI | *p* |
| Intercept | 848.34 | 790.72 to 905.95 | **<0.001** | 2.65 | 1.30 to 4.00 | **<0.001** |
| Loss aversion | 25.37 | 10.96 to 39.77 | **<0.001** | 4.00 | 2.69 to 5.31 | **<0.001** |
| Betrayal aversion | -35.35 | -65.84 to -4.85 | **0.023** | 8.66 | 6.29 to 11.04 | **<0.001** |
| Observations | 9787 |  |  | 9787 |  |  |

*Notes*: The final model for Decision times as: Decision times ~ Loss aversion + Betrayal aversion + (Betrayal aversion | Participant), and for the proportion accepted as: proportion accepted ~ Loss aversion + Betrayal aversion + (Loss aversion + Betrayal aversion | Participant). Loss aversion and betrayal aversion were standardized before being entered into the model. Statistically significant *p* values (<0.05, two-sided) are shown in bold. CI = confidence interval.

**Table S5.** Results of linear mixed-effects models predicting P3 (left) and LPP (right) as a function of loss aversion and betrayal aversion. (*N*=49)

|  | P3 |  |  | LPP |  |  |
|---|---|---|---|---|---|---|
| Predictors | *Estimates* | 95% CI | *p* | *Estimates* | 95% CI | *p* |
| Intercept | 3.88 | 3.19 to 4.58 | **<0.001** | 2.54 | 1.86 to 3.22 | **<0.001** |
| Loss aversion | -0.07 | -0.32 to 0.18 | 0.566 | -0.33 | -0.57 to -0.08 | **0.009** |
| Betrayal aversion | 0.45 | 0.09 to 0.81 | **0.015** | 0.68 | 0.32 to 1.04 | **<0.001** |
| Observations | 9425 |  |  | 9425 |  |  |

*Notes*: The final model for P3/ LPP as: P3 amplitude/ LPP amplitude ~ Loss aversion + Betrayal aversion + (Loss aversion \| Participant). Loss aversion and betrayal aversion were standardized before being entered into the model. Statistically significant *p* values (<0.05, two-sided) are shown in bold. CI = confidence interval.

**Table S6.**
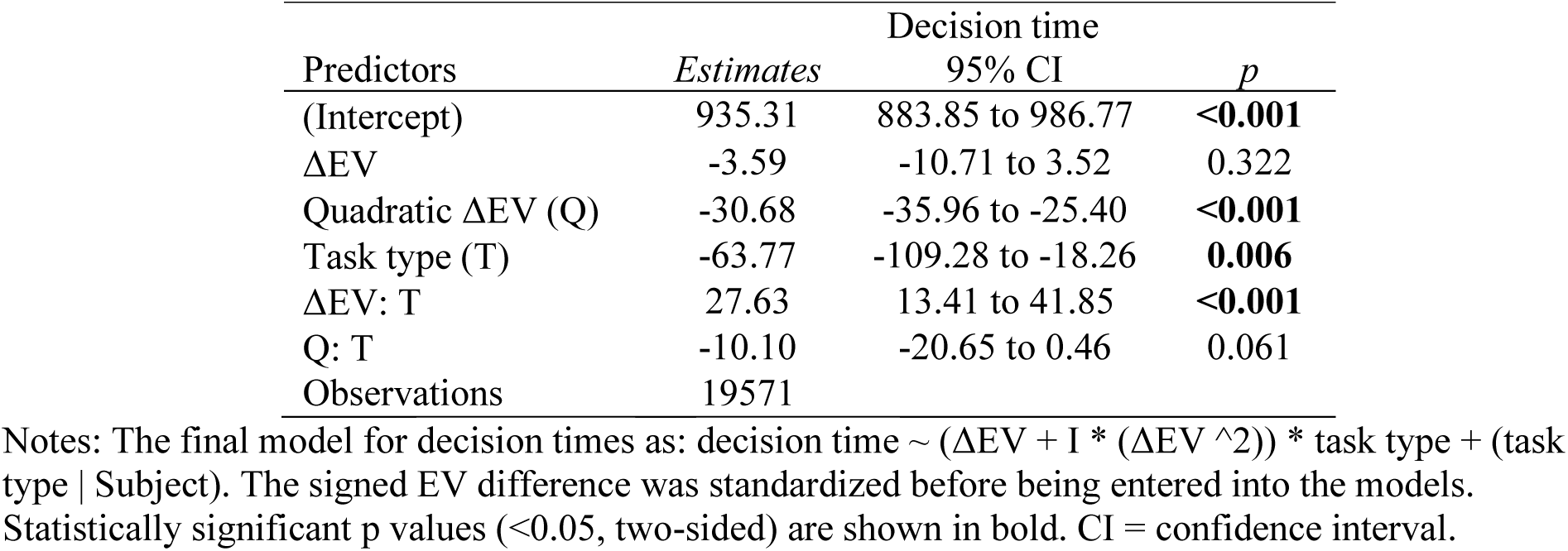
Results of linear mixed-effects models predicting decision time as a function of ΔEV, Quadratic ΔEV and task type. (*N*=49) Decision time.

| Predictors | Decision time |  |  |
|---|---|---|---|
|  | *Estimates* | 95% CI | *p* |
| (Intercept) | 935.31 | 883.85 to 986.77 | **<0.001** |
| ΔEV | -3.59 | -10.71 to 3.52 | 0.322 |
| Quadratic ΔEV (Q) | -30.68 | -35.96 to -25.40 | **<0.001** |
| Task type (T) | -63.77 | -109.28 to -18.26 | **0.006** |
| ΔEV: T | 27.63 | 13.41 to 41.85 | **<0.001** |
| Q: T | -10.10 | -20.65 to 0.46 | 0.061 |
| Observations | 19571 |  |  |

Notes: The final model for decision times as: decision time ~ (ΔEV + I \* (ΔEV ^2)) \* task type + (task type | Subject). The signed EV difference was standardized before being entered into the models. Statistically significant p values (<0.05, two-sided) are shown in bold. CI = confidence interval.

**Table S7.** Results of the generalized linear mixed-effects models predicting choices as a function of probability, principal, and task type, before (left) and after (right) participant exclusions.

|  | Choices (original; N=49) |  |  | Choices (after exclusions; N=41) |  |  |
|---|---|---|---|---|---|---|
| Predictors | *Log odds* | 95% CI | *p* | *Log odds* | 95% CI | *p* |
| Intercept | 0.58 | 0.30 to 0.86 | <0.001 | 0.55 | 0.26 to 0.84 | **<0.001** |
| Probability (Pro) | 3.00 | 2.70 to 3.31 | <0.001 | 3.04 | 2.70 to 3.37 | **<0.001** |
| Principal (Pri) | -0.42 | -0.47 to -0.37 | <0.001 | -0.49 | -0.55 to -0.44 | **<0.001** |
| Task type (T) | -0.83 | -1.08 to -0.57 | <0.001 | -0.92 | -1.19 to -0.65 | **<0.001** |
| Pro:Pri | 0.31 | 0.25 to 0.37 | <0.001 | 0.36 | 0.29 to 0.43 | **<0.001** |
| Pro:T | -0.44 | -0.58 to -0.30 | <0.001 | -0.52 | -0.67 to -0.36 | **<0.001** |
| Pri:T | -0.10 | -0.20 to -0.01 | 0.029 | -0.12 | -0.22 to -0.01 | **0.029** |
| Pro: Pri: T | -0.20 | -0.32 to -0.07 | 0.002 | -0.26 | -0.40 to -0.12 | **<0.001** |
| Observations | 19571 |  |  | 16378 |  |  |

*Notes*: The same model was fitted before and after participant exclusions: Proportion accepted ~ Probability \* Principal \* Task type+ (Probability + Task type \| Participant). Probability and principal level were standardized before being entered into the model. Statistically significant *p* values (<0.05, two-sided) are shown in bold. CI = confidence interval.

**Table S8.** Results of the linear mixed-effects models predicting decision times as a function of probability, principal, and task type, before (left) and after (right) participant exclusions.

|  | Decision times (original; N=49) |  |  | Decision times (after exclusions; N=41) |  |  |
|---|---|---|---|---|---|---|
| Predictors | *Estimates* | 95% CI | *p* | *Estimates* | 95% CI | *p* |
| Intercept | 904.64 | 853.45 to 955.84 | **<0.001** | 897.35 | 843.80 to 950.90 | **<0.001** |
| Probability (Pro) | -21.51 | -28.05 to -14.96 | **<0.001** | -21.01 | -28.23 to -13.80 | **<0.001** |
| Principal (Pri) | -26.02 | -32.57 to -19.48 | **<0.001** | -27.27 | -34.48 to -20.05 | **<0.001** |
| Task type (T) | -73.89 | -118.16 to -29.63 | **0.002** | -77.38 | -126.90 to -27.87 | **0.002** |
| Pro:Pri | 12.97 | 6.42 to 19.51 | **<0.001** | 14.48 | 7.26 to 21.70 | **<0.001** |
| Pro:T | 26.55 | 13.45 to 39.64 | **<0.001** | 28.24 | 13.81 to 42.67 | **<0.001** |
| Pri:T | -13.33 | -26.42 to -0.24 | **0.046** | -9.96 | -24.39 to 4.47 | 0.176 |
| Pro: Pri: T | 5.16 | -7.93 to 18.25 | 0.440 | -2.13 | -16.56 to 12.31 | 0.773 |
| Observations | 19571 |  |  | 16378 |  |  |

*Notes*: The same model was fitted before and after participant exclusions: Decision times ~ Probability \* Principal \* Task type+ (Task type \| Participant). Probability and principal level were standardized before being entered into the model. Statistically significant *p* values (<0.05, two-sided) are shown in bold. CI = confidence interval.

**Table S9.**
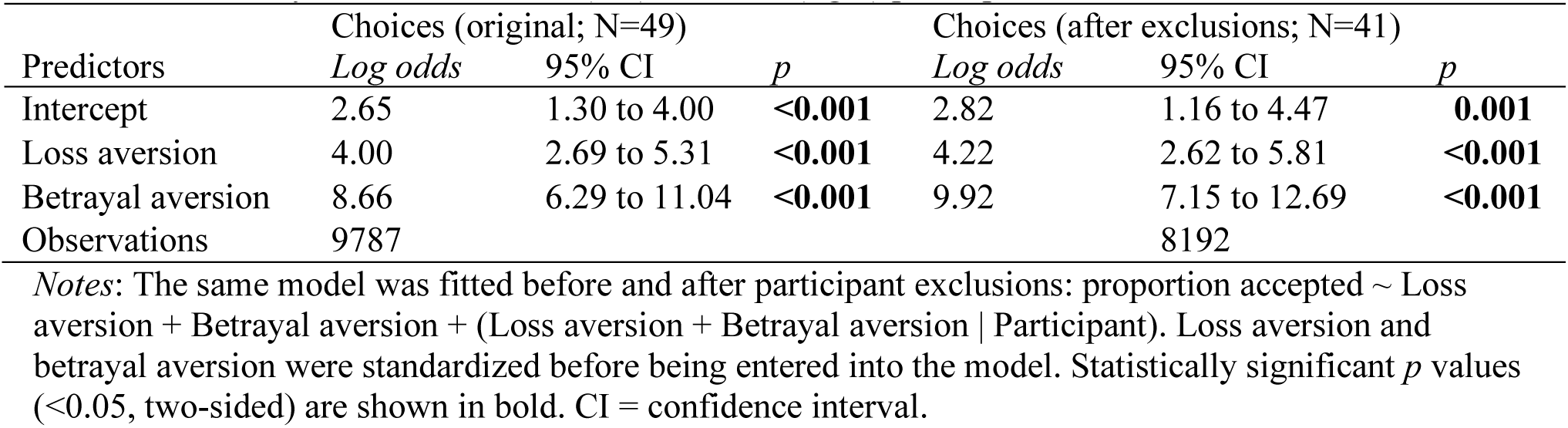
Results of the generalized linear mixed-effects models predicting choices as a function of loss aversion and betrayal aversion, before (left) and after (right) participant exclusions.

**Table S10.** Results of the linear mixed-effects models predicting decision times as a function of loss aversion and betrayal aversion, before (left) and after (right) participant exclusions.

|  | Decision times (original; N=49) |  |  | Decision times (after exclusions; N=41) |  |  |
|---|---|---|---|---|---|---|
| Predictors | *Estimates* | 95% CI | *p* | *Estimates* | 95% CI | *p* |
| Intercept | 848.34 | 790.72 to 905.95 | **<0.001** | 836.59 | 774.41 – 898.78 | **<0.001** |
| Loss aversion | 25.37 | 10.96 to 39.77 | **<0.001** | 26.33 | 10.31 – 42.34 | **0.001** |
| Betrayal aversion | -35.35 | -65.84 to -4.85 | **0.023** | -37.26 | -75.54 – 1.02 | 0.056 |
| Observations | 9787 |  |  | 8192 |  |  |

*Notes*: The same model was fitted before and after participant exclusions: Decision times ~ Loss aversion + Betrayal aversion + (Betrayal aversion | Participant). Loss aversion and betrayal aversion were standardized before being entered into the model. Statistically significant *p* values (<0.05, two-sided) are shown in bold. CI = confidence interval.

**Table S11.** Results of linear mixed-effects models predicting P3 as a function of loss aversion and betrayal aversion, before (left) and after (right) participant exclusions.

|  | P3 (original; N=49) |  |  | P3 (after exclusions; N=41) |  |  |
|---|---|---|---|---|---|---|
| Predictors | *Estimates* | 95% CI | *p* | *Estimates* | 95% CI | *p* |
| Intercept | 3.88 | 3.19 to 4.58 | **<0.001** | 4.20 | 3.40 – 5.00 | **<0.001** |
| Loss aversion | -0.07 | -0.32 to 0.18 | 0.566 | -0.15 | -0.45 – 0.14 | 0.300 |
| Betrayal aversion | 0.45 | 0.09 to 0.81 | **0.015** | 0.58 | 0.14 – 1.02 | **0.009** |
| Observations | 9425 |  |  | 7893 |  |  |

*Notes*: The final model for P3 as: P3 amplitude/ LPP amplitude ~ Loss aversion + Betrayal aversion + (Loss aversion \| Participant). Loss aversion and betrayal aversion were standardized before being entered into the model. Statistically significant *p* values (<0.05, two-sided) are shown in bold. CI = confidence interval.

**Table S12.**
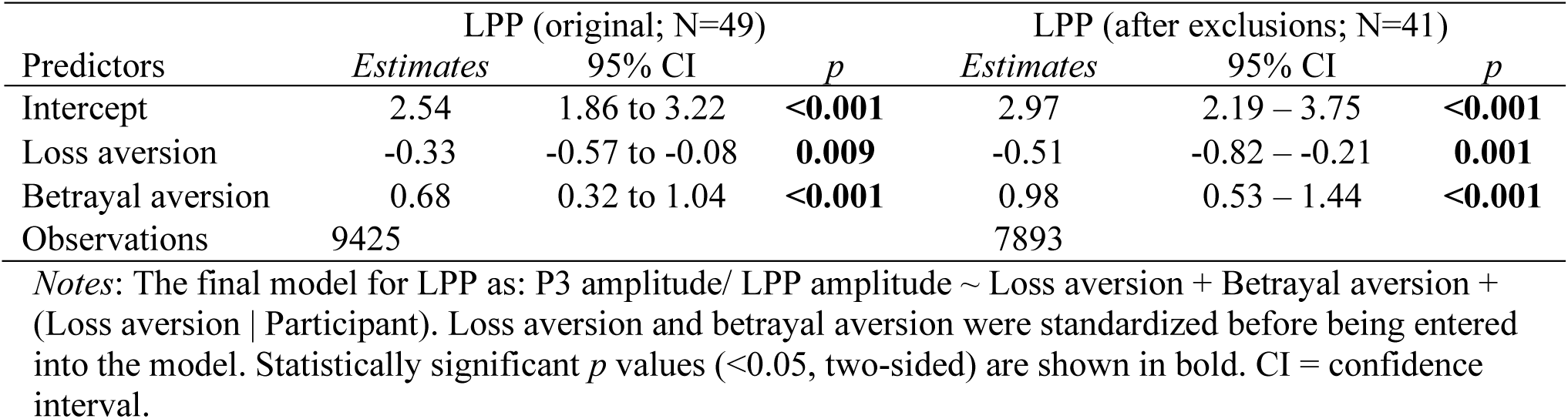
Results of linear mixed-effects models predicting LPP as a function of loss aversion and betrayal aversion, before (left) and after (right) participant exclusions.

|  | LPP (original; N=49) |  |  | LPP (after exclusions; N=41) |  |  |
|---|---|---|---|---|---|---|
| Predictors | *Estimates* | 95% CI | *p* | *Estimates* | 95% CI | *p* |
| Intercept | 2.54 | 1.86 to 3.22 | **<0.001** | 2.97 | 2.19 – 3.75 | **<0.001** |
| Loss aversion | -0.33 | -0.57 to -0.08 | **0.009** | -0.51 | -0.82 – -0.21 | **0.001** |
| Betrayal aversion | 0.68 | 0.32 to 1.04 | **<0.001** | 0.98 | 0.53 – 1.44 | **<0.001** |
| Observations | 9425 |  |  | 7893 |  |  |

*Notes*: The final model for LPP as: P3 amplitude/ LPP amplitude ~ Loss aversion + Betrayal aversion + (Loss aversion \| Participant). Loss aversion and betrayal aversion were standardized before being entered into the model. Statistically significant *p* values (<0.05, two-sided) are shown in bold. CI = confidence interval.

**Figure S1.**
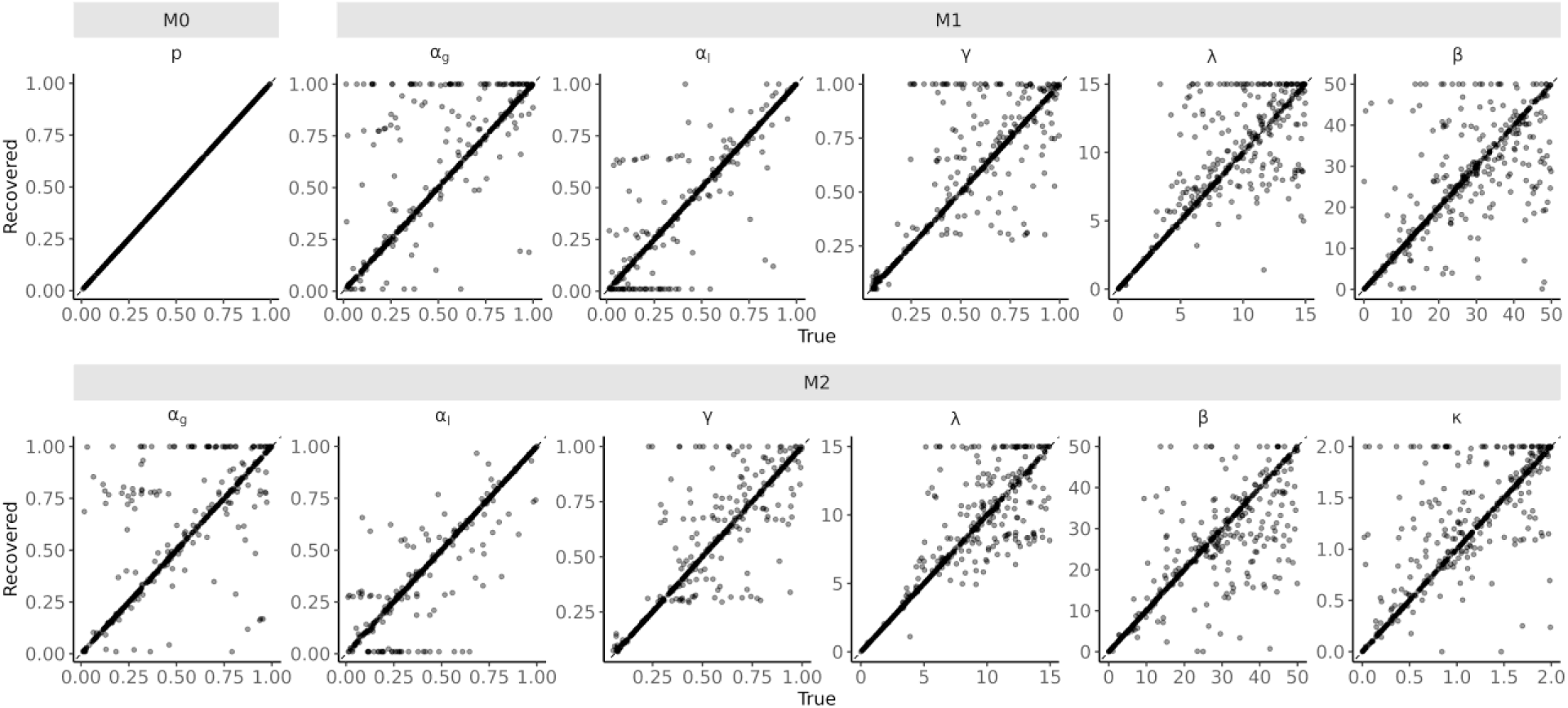
The scatter plot of parameter recovery analysis of Model 0 (A), Model 1 (B) and Model 2 (C).

**Figure S2.**
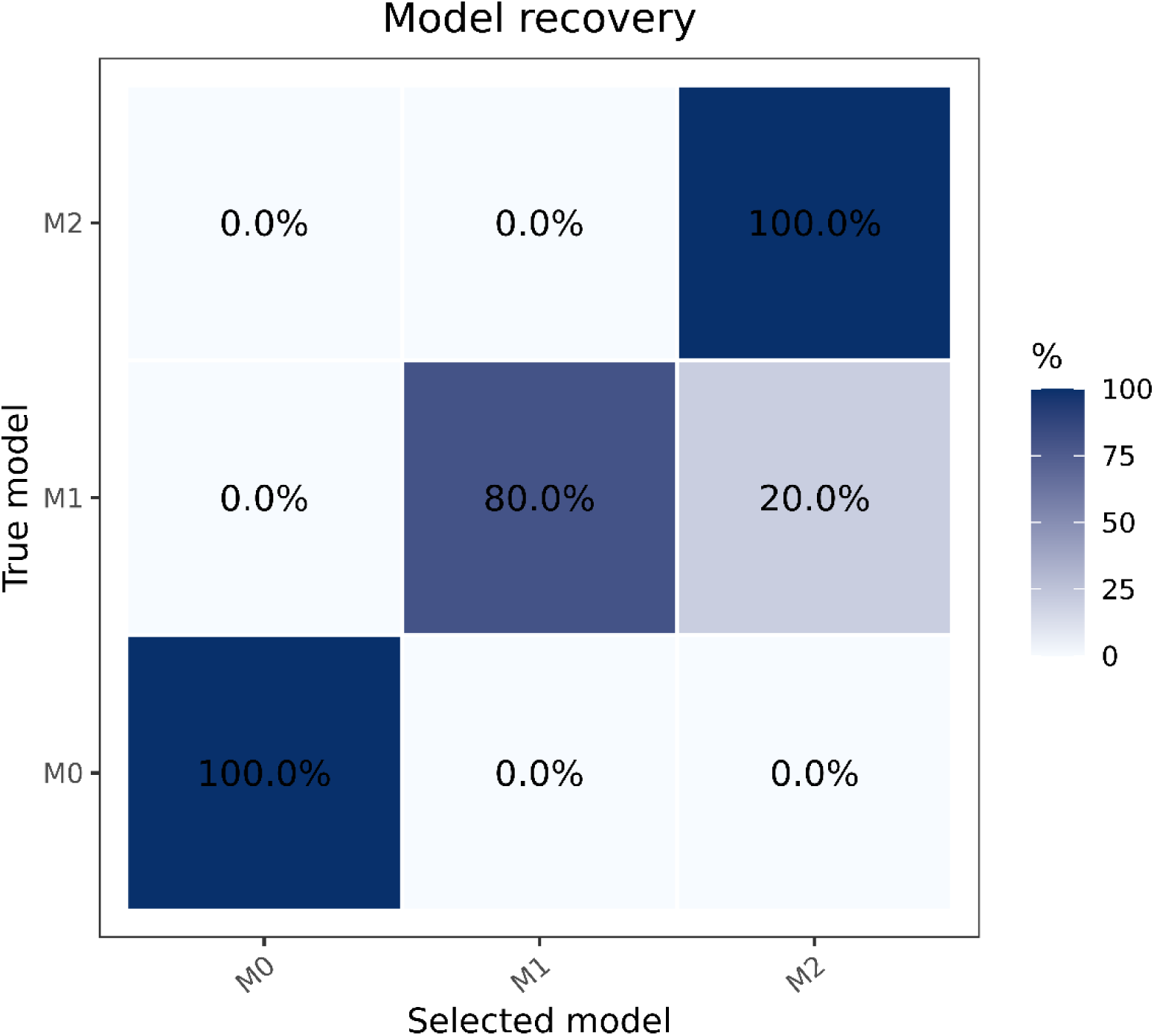
The confusion matrix of model recovery analysis.

**Figure S3.**
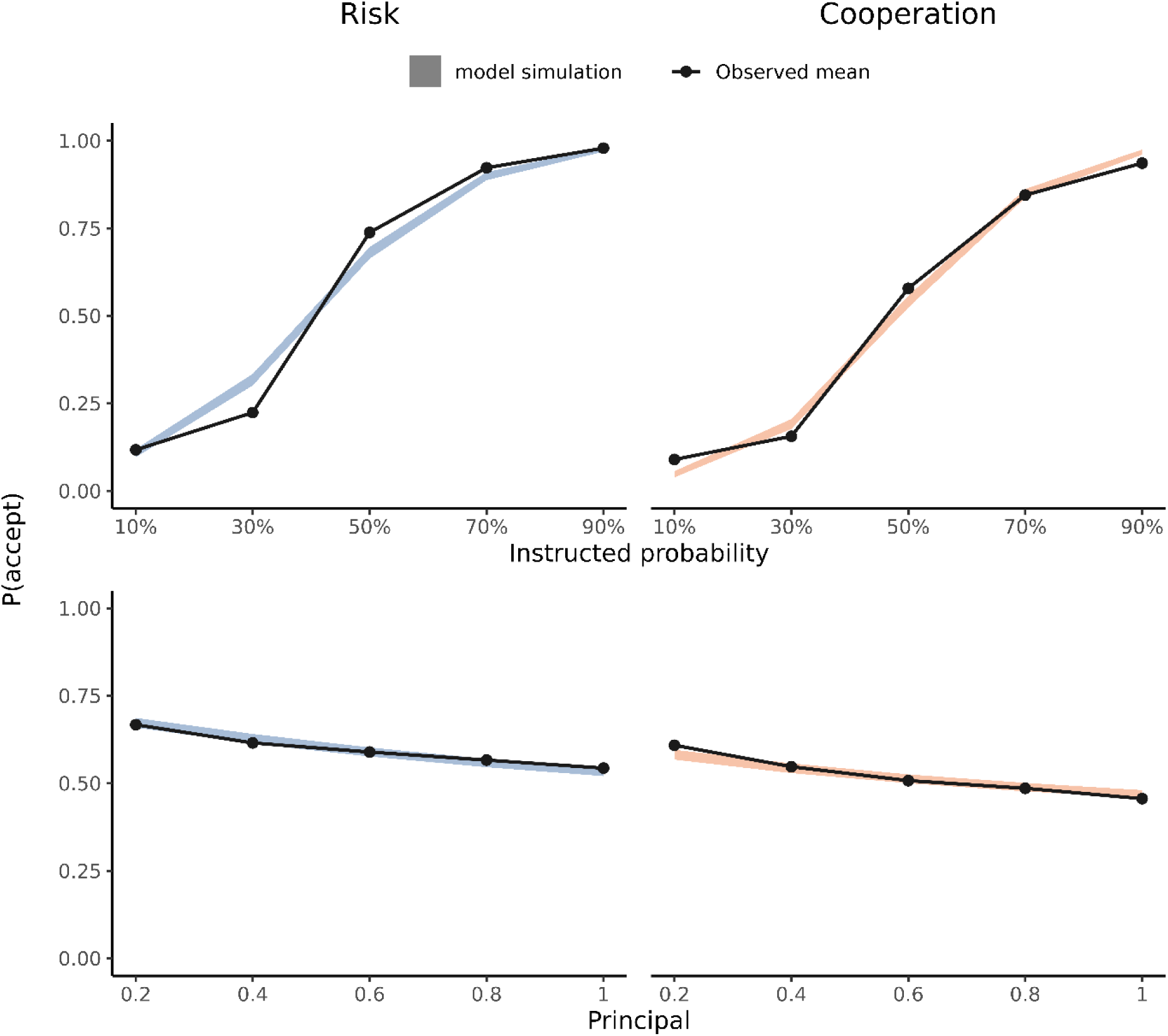
Model-based simulations as a function of probability level (top) and principal magnitude (bottom). Shaded areas show simulated values; black lines with points show observed means.

